# Size-Modulated Mesoderm-Endoderm Divergence and Myocardial Cavitation in Micropatterned Cardioids

**DOI:** 10.1101/2025.09.17.676874

**Authors:** Plansky Hoang, David W. McKellar, Andrew Kowalczewski, Nhu Y. Mai, Meng Chai, Xiaojun L. Lian, Yi Zheng, Jeffrey Amack, Nathan Tucker, Iwijn De Vlaminck, Huaxiao Yang, Benjamin D. Cosgrove, Zhen Ma

## Abstract

The human heart, originating from the splanchnic mesoderm, is the first functional organ to develop, co-evolving with the foregut endoderm through reciprocal signaling. Previously, cardioid models offered new insights on cardiovascular cell lineages and tissue morphogenesis during heart development, while mesoderm-endoderm crosstalk remain incompletely understood. Here, we integrated micropatterned cardioids, CRISPR-engineered reporter hiPSCs, deep-tissue imaging, and single-cell RNA sequencing (scRNA-seq) to explore synergistic mesoderm-endoderm co-development. scRNA-seq with PHATE trajectory mapping reconstructed lineage bifurcations of mesoderm-heart and endoderm-foregut lineages, identifying key cell types in cardiac and hepatic development. Ligand-receptor interaction analysis highlighted mesodermal cells enriched in non-canonical WNT, NRG, and TGF-β signaling, while endodermal cells exhibited VEGF and Hedgehog activity. We found that micropattern sizes influenced cellular composition, cardioid cavitation, contractile functions, and mesoderm-endoderm signaling crosstalk. The cardioids generated from 600 µm diameter circle patterns showed larger cavity formation resembling early heart chamber formation. Our findings establish micropatterned cardioids as a model for mesoderm-endoderm co-development, enhancing our understanding of heart-foregut synergy during early embryogenesis.

## INTRODUCTION

The human heart originates from the splanchnic mesoderm, as the first functional organ to form during mammalian development. Its proper development not only establishes the foundation for the development of nearby organs but also relies on signaling from neighboring tissues. During embryogenesis, endodermal gut tube expands in parallel with the co-developing heart tube^1^. Directional rearrangement of cardiac cells during heart tube extension is coordinated with accompanying foregut endoderm at the midline^2,3^. Cell fate decisions in the co-emergence of the heart and liver rely on inductive paracrine signaling between heart-forming region and developing foregut endoderm^4–7^. Studies utilizing tissue explants revealed that the growth factors released by the endodermal cells promote cardiac development and heart formation^8^. At the same time, hepatic specification requires the presence of the surrounding cardiac mesoderm, septum transversum, and endothelium^9^. These findings underscore the importance of reciprocal influence and cellular crosstalk between the co-developing mesoderm-heart and endoderm-foregut lineages.

Stem cell-derived organoids are designed to mimic early developing organs by allowing differentiating cells to self-organize into functional tissue structures. Recent advancements in heart-forming organoids, also called “cardioids”, have shown promising results in replicating appropriate lineage specification and key morphogenetic events during heart development. By precisely coordinating WNT, FGF and BMP4 signaling, cardioids generated from human or murine pluripotent stem cells (PSCs) modeled fetal heart development, encompassing all major cardiovascular cell lineages^10–12^. These cardioids exhibited chamber-like cavity formation, resembling the natural structure of the developing heart^13,14^. Notably, the presence of endodermal cells within the cardioids highlighted the co-development of cardiac mesoderm and foregut endoderm. In a gastruloid model, the heart-forming region emerged adjacent to the primitive gut-like and endocardial-like layers^15^. In cardioid models specifically, co-development of cardiac tissues in conjunction with endodermal derivatives played a critical role in spatial tissue morphogenesis and myocardial maturation^16,17^. However, most studies to date have focused on individual lineages or signaling pathways, and lack a comprehensive understanding of the temporally dynamic, combinatorial signaling network in mesoderm-endoderm synergy. The precise lineage divergence of mesoderm-endoderm within cardioids has not been well established, underscoring the need for further research into their complex interplay during heart-foregut development.

Micropatterning hPSCs provides a powerful platform to study embryogenesis and organogenesis by enabling precise spatial control over cell adhesion, geometry, and signaling environments. By constraining hPSCs into defined patterns, researchers can recapitulate key symmetry-breaking events, germ layer specification, and morphogen gradients that drive early embryonic organization^18–23^. Previously, we established a cardioid model by micropatterning human induced pluripotent stem cells (hiPSCs)^24,25^. In this study, we employed CRISPR-engineered iPSC reporter lines, advanced tissue imaging techniques, and single-cell RNA sequencing (scRNA-seq) to elucidate the mesoderm-endoderm co-emergence within our cardioids. By generating cardioids from 200 µm, 600 µm and 1000 µm diameter circles, we found that pattern sizes altered multi-cellular composition, modulated cardioid internal cavitation, and influenced their contractile functions. Ligand-receptor interaction analysis demonstrated that mesoderm-derived cells primarily contributed to non-canonical WNT, NRG, and TGF-β signaling, while endoderm-derived cells were enriched in VEGF and Hedgehog signaling, highlighting the potential for modeling heart development and providing mechanistic understanding of heart-foregut crosstalk.

## RESULTS

### Multicellular Co-Developing Cardioids with Nascent Myocardial Cavities

We micropatterned hiPSC colonies into 600 µm diameter circles to generate cardioids using the WNT-modulation protocol (**Figure 1a**). We fixed, stained, and imaged the whole-mounted cardioids to visualize their intact structure without tissue sectioning (**Movie S1**). We identified cardiomyocytes at the dome region by staining ACTN2 (sarcomeric α-actinin), TNNI2 (cardiac troponin I), and MYH6 (α-myosin heavy chain), and stromal cells at the peripheral region by staining for TAGLN (SM22), VIM (vimentin), CNN1 (calponin), and ACTA2 (smooth muscle actin) (**Figure S1**). By reconstructing the Z-stack of confocal images, we were able to visualize the dome structure of these micropatterned cardioids from an orthogonal view. However, due to the limited depth of light penetration in confocal microscopy, we were unable to visualize the cardioid internal structure.

**Figure 1.**
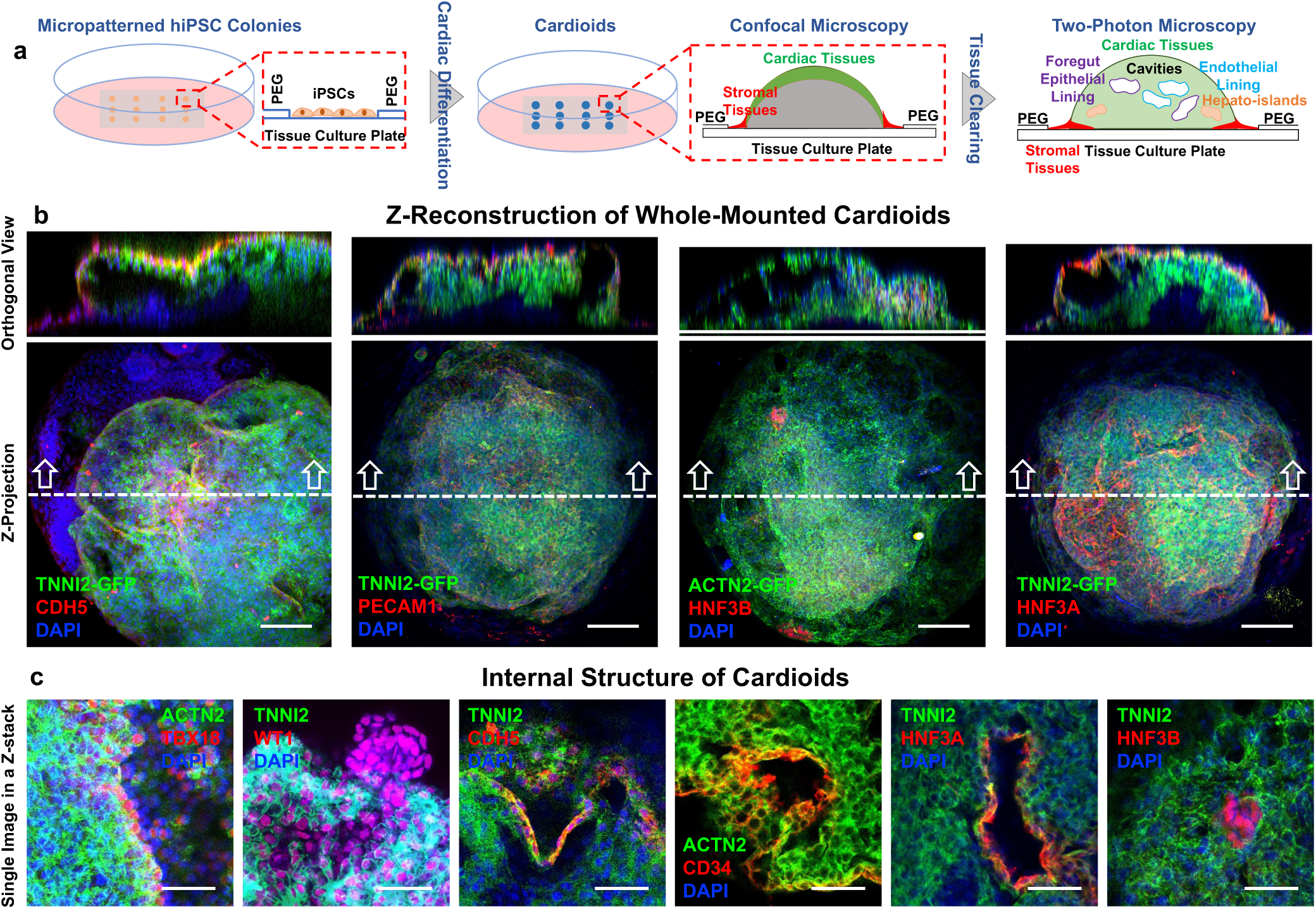
Deep-tissue imaging reveals mesoderm and endoderm cell types in micropatterned cardioids. (a) Schematic representation of cardioid generation from micropatterned hiPSCs. Whole-mounted cardioids were treated with tissue-clearing reagents to enable deep-tissue imaging. (b) Representative 3D reconstruction images of two-photon microscopy of micropatterned cardioids derived from TNNI2-GFP and ACTN2-GFP hiPSC reporter lines. Cardioids were stained with endothelial markers CDH5 (VE-cadherin) and PECAM1 (CD31) for vascular structures and HNF3A and HNF3B for foregut cells. Scale: 100 µm. (c) Representative images of internal structure of TNNI2-GFP and ACTN2-GFP cardioids with epicardial markers (TBX18 and WT1), endothelial markers (CDH5 and PECAM1), and foregut markers (HNF3A and HNF3B). These representative images from different cardioids reveals endothelial and foregut epithelial cells lining the cavities within the cardioids. Scale: 50 µm.

For deep tissue imaging inside the cardioids, we used two CRISPR-engineered hiPSC lines with fluorescent reporters for cardiomyocytes (TNNI-GFP and ACTN-GFP) to generate the cardioids, which were then optically cleared prior to immunostaining. We imaged the whole-mounted and intact cardioids using two-photon microscopy for deep tissue penetration (**Figure 1b, Movie S2**). Z-projection of the resulting 3D images revealed a prominent population of cardiomyocytes, along with staining for foregut endodermal cells by HNF3A and HNF3B and endothelial cells by PECAM1 (CD31) and CDH5 (VE-cadherin). Notably, the orthogonal view provided a clear visualization of the internal structure, including the cavity formation within the cardioids. Though tissue clearance enhanced the quality of deep tissue imaging, total volume of the cardioids was reduced due to the clearance procedures, compared to the confocal images (**Figure S1**). By selecting single images from the Z-stacks, we were able to visualize the cardioid cavities (**Figure 1c**). The epicardial cells (WT1 and TBX18) were predominantly localized at the outer layer of the cardioids. Both endothelial cells (CDH5 and PECAM1) and foregut epithelial cells (HNF3α) lined the cavities inside the cardioids. In contrast, HNF3β+ liver progenitors were found to be clustered in small regions inside the cardioids, indicating the emergence of endoderm islands. Previous studies of HFOs also demonstrated two main phenotypic structures of cavity-lining foregut endoderm lumens and rosette-like foregut endoderm islands, similar to our observations^16^.

### Mesoderm-Endoderm Divergence in PHATE Trajectory

To confirm the emergence of endoderm lineage observed by immunostaining in micropatterned cardioids, we performed scRNA-seq (Chromium v3.1, 10X Genomics) on 600µm micropatterned samples at seven timepoints (Day 0, 1, 4, 6, 8, 12, 21) spanning cardioid differentiation. We additionally assayed the samples generated with 200 µm and 1000 µm micropatterned at Day 4 and Day 21 (**Figure 2a, Figure S2 & S3**), as Day 4 corresponds to the early mesoderm–endoderm divergence and Day 21 represents the terminal stage of cardioid characterization. Following data preprocessing and quality control, we retained 96,401 cells for subsequent transcriptome analysis (**Figure S2a**). We first integrated the scRNA-seq data using Harmony to remove both batch effects and cell cycle signatures^26^ (**Figure S2b**). Next, we applied PHATE (Potential of Heat-diffusion for Affinity-based Transition Embedding), a dimensional reduction algorithm which is well-suited for data with continuous and branching latent structure for visualization of mesoderm-endoderm divergence^27^ (**Figure 2b-2f**). Finally, we annotated cell clusters based on canonical gene markers (**Figure 2g**). We categorized the cells into 13 distinct types, which not only provided insights into the multicellular composition of the cardioids but also reconstructed the developmental divergence of mesoderm-heart and endoderm-liver lineages during cardioid development.

**Figure 2.**
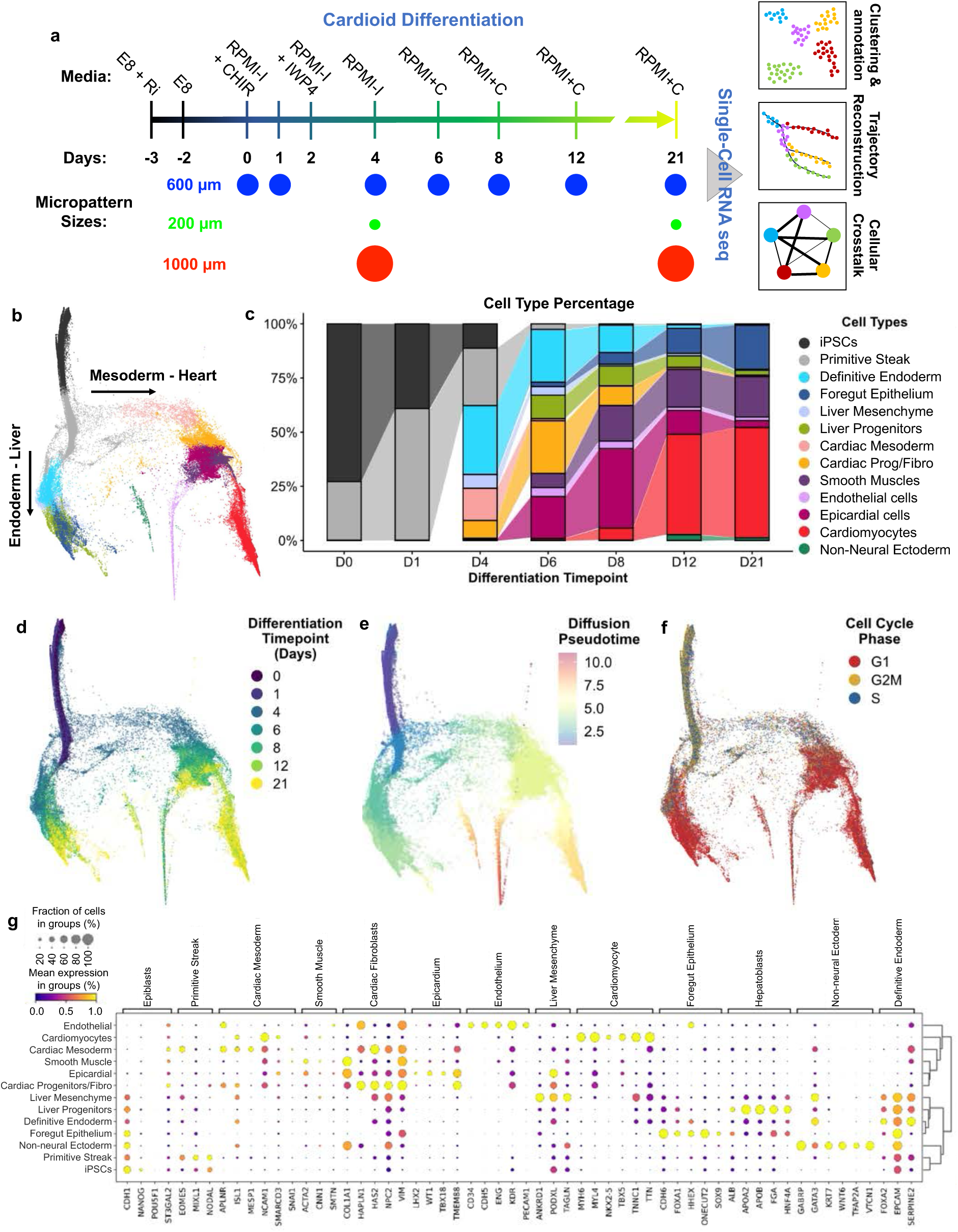
Single-cell transcriptomics reveals the divergence of mesoderm-heart and endoderm-liver lineages. (a) Schematic of the cardioid differentiation protocol and sample collection for scRNA-seq analysis. Seven samples were collected from 600 µm micropatterned cardioids at Day 0, 1, 4, 6, 8, 12, and 21, while two samples were collected from 200 µm and 1000 µm patterns at Day 4 and 21. (b) PHATE embedding illustrates the developmental bifurcation of the mesoderm-heart and endoderm-liver lineages. (c) Visualization of cell fate transitions throughout cardioid differentiation. Illustrating (d) differentiation time points, (e) pseudo-time progression, and (f) cell cycle states on PHATE embedding. (g) Cell type annotations based on canonical gene expression markers.

At Day 0, two cell types were identified from the confluent micropatterned cell colonies, iPSCs (*NANOG*, *POU5F1*, *CDH1*) and primitive streak (PS) cells (*MIXL1*, *NODAL*, *EOMES*), suggesting that geometric confinement to densely packed hiPSCs triggered the spontaneous differentiation into the primitive streak (PS) cells. Upon WNT activation on Day 1, iPSCs underwent rapid differentiation into the PS cells (**Figure 2c**). Early mesoderm-endoderm divergence was observed at Day 4 after WNT inhibition through IWP4 treatment (**Figure 2d**). At this stage, both cardiac mesoderm (*MESP1*, *ISL1*, *APLNR*) and definitive endoderm (*EPCAM*, *FOXA2*, *SERPINE2*) co-emerged within the cardioids. Surprisingly, starting from Day 8, a non-neural ectoderm (*KRT7*, *WNT6*, *TFAP2A*, and *GABRP*) cell cluster emerged between mesoderm-heart and endoderm-liver lineages (**Figure 2e**). Non-neural ectoderm cells are generally considered the cells that give rise to the epidermis and the amniotic membrane, but not nervous tissues^28–30^. This result illustrates the co-development of all three germ layers in the micropatterned cardioids, which has not been reported previously.

We identified 6 clusters in the mesoderm-heart lineage and 4 cell clusters in the endoderm-foregut lineage (**Figure 2g**). In the mesoderm lineage, cardiac mesoderm developed into cardiac progenitors/fibroblasts (*NPC2*, *VIM*), epicardial cells (*WT1, TBX18*), and smooth muscle cells (*ACTA2* and *CNN1*). Next, the mesoderm lineage further bifurcated into endocardial cells (*CD34*, *PECAM1*, *CDH5*, *KDR*) and cardiomyocytes (*TTN*, *MYH6*, *MYL4*, *NKX2-5*, *ISL1).* In the endoderm lineage, definitive endoderm developed into liver progenitors (*ALB*, *APOA2*, and *HNF4A*), foregut epithelial cells (*CDH6*, *FOXA1*, and *ONECUT2*), and liver mesenchyme (*PODXL*, *ANKRD1*, and *TAGLN*). Liver mesenchymal cells expressed both endoderm-epithelial markers (*GATA3*, *FOXA2*, and *EPCAM*) and mesoderm-mesenchymal markers (*PODXL*, *TAGLN*, *NPC2*, and *VIM*), indicating their gene expression pattern is extensively influenced by the mesoderm lineage.

### Lineage-Specific Multicellular Development

The developmental trajectory of mesoderm-endoderm divergence was recapitulated in the PHATE embedding. To better resolve cell composition in each lineage, we performed PAGA (Partition-based Graph Abstraction)-guided force-directed dimensional reduction for sub-clustering mesoderm lineage and endoderm lineage separately (**Figure 3**). PAGA clarified the relationships between the mesoderm cells (**Figure 3a**) and aided further sub-clustering the mesoderm cells into 10 populations (**Figure 3b).** Cardiac mesoderm cells (*EOMES, APLNR, MESP1*) were progressed into two cardiac progenitor clusters. Early progenitors (Cardiac Progenitor 2: PPRX1+/BMPER^high^) were developmentally closer to the cardiac mesoderm but bifurcated into both cardiomyocytes and endothelial cells. Late progenitors (Cardiac Progenitor 1: LHX2+/PITX2^high^) were more committed to the cardiac stromal lineages, including fibroblasts (POSTN, CXCL12, NR2F2), epicardial cells (WT1, TBX18, SOX6), and smooth muscle cells (TAGLN, CCN2, and ANXA2) (**Figure 3c**)^31,32^. We further constructed the pseudo-time for three cardiomyocyte clusters using CellRank^33^ (**Figure S4a**) and analyzed a comprehensive list of cardiomyocyte-related genes base on a previous report on human heart development^34^. Genes involved in early cardiac transcription (PITX2, HMGA1, ISL1) and signaling molecules (FGFs and BMPs) were highly expressed in early cardiomyocytes (Cardiomyocytes 3) (**Figure S4a**). We observed that transcription factors (HEY2 and IRX4), sarcomere protein (LDB3, MYL3), potassium ion channels (KCNH2, KCNH7), and calcium handling (TRDN, TYR2, CMYA5) progress towards the end of pseudo-time (**Figure S4b**). A similar maturation progress was observed across differentiation time points. Day 6/8 cardiomyocytes exhibited higher expression of signaling molecules, whereas Day 21 cardiomyocytes showed increased gene expression of sarcomeres and calcium handling (**Figure S4b**). For Day 21 cardiomyocytes, 200 µm cardioids exhibited higher expression of transcription factors, while 600 µm cardioids displayed elevated expression of sarcomere and calcium handling genes (**Figure S4c**). These findings suggest that cardiomyocytes from 600 µm cardioids exhibit a higher maturation level compared to those from other cardioids.

**Figure 3.**
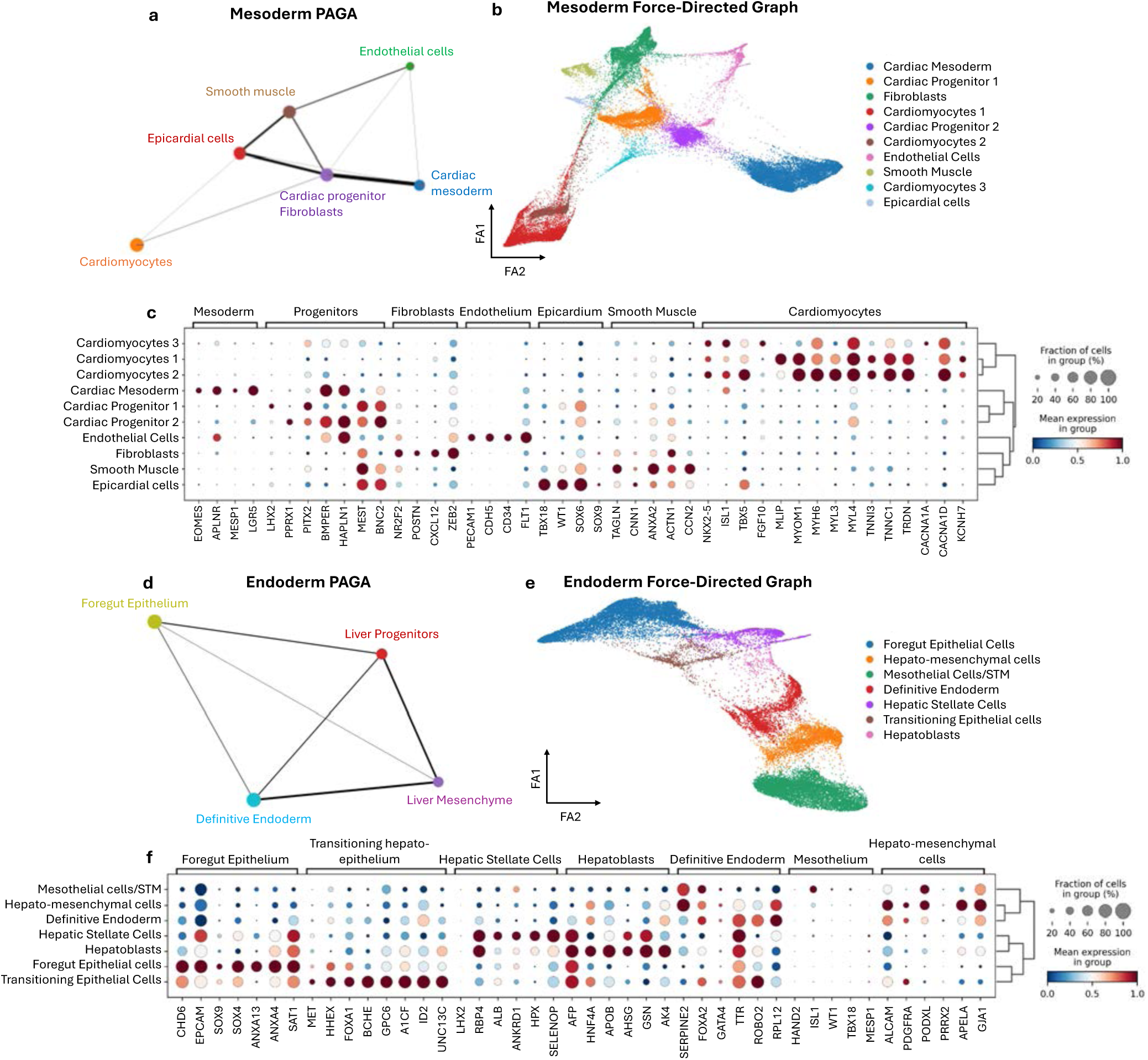
Force-directed sub-clustering of mesoderm-heart and endoderm-liver lineages. (a) Partition-based graph abstraction (PAGA) was extracted from PHATE embedding and used to guide (b) force-directed dimensional reduction of mesoderm lineage. Cells progressed from cardiac mesoderm to cardiac progenitors, followed by bifurcation into cardiomyocytes, endothelial cells, and stromal cells (fibroblasts, smooth muscle cells, and epicardial cells). (c) Canonical gene expression was profiled for mesodermal cell clusters. (d) PAGA and (e) force-directed graph of endoderm lineage further delineated liver mesenchyme into mesoderm-origin mesothelial cells, endoderm-origin hepato-mesenchymal cells, and hepatic stellate cells. (c) Canonical gene expression was profiled for endodermal cell clusters.

A similar PAGA-guided force-directed process was also applied to further sub-cluster the endoderm lineage into 7 cell populations (**Figure 3d and 3e**), where definitive endoderm (FOXA2, SERPINE2, TTR), foregut epithelial cells (EPCAM, SOX9, SOX4), and hepatoblasts (AFP, HNF4A, ALB) were better separated. Importantly, liver mesenchyme was further divided into three cell sub-types from different developmental origins (**Figure 3f**): hepato-mesenchymal cells, mesothelial cells, and hepatic stellate cells, which is consistent with the recent scRNAseq studies on liver mesenchyme development^35^. The mesothelial cells (*WT1, TBX18*) are derived from STM that separates the heart and liver during embryonic development^36,37^. Their mesoderm origin was confirmed by the expression of ISL1, HAND2, and MESP1^38,39^. In contrast, hepato-mesenchymal cells (APOA1, HNF4A) originate from the endoderm but undergo epithelial-mesenchymal transition (ALCAM, PODXL, PDGFRA) as hepato-epithelial cells migrate into the STM^35,40^. Notably, a small cluster of hepatic stellate cells (LHX2) was identified, which share gene expression patterns with hepatoblasts (RBP4, ALB, AFP) but also express fibrogenesis-associated genes (HPX, ANKRD1)^41,42^. Developmentally, the liver bud contains a great diversity of mesodermal cell states, likely due to the unique process in which hepatic endoderm delaminates and invades the adjacent STM. This invasion requires more intricate epithelial-mesenchymal interactions compared to other gastrointestinal organs, which primarily form through epithelial evagination^39^.

### The Impact of Pattern Sizes on Mesoderm-Endoderm Divergence

To visualize mesoderm-endoderm divergence, we micropatterned and differentiated human embryonic stem cells (RUES2-GLR) expressing triple reporters for SOX2 (GFP, epiblast), TBXT (mCer, primitive streak), and SOX17 (RFP, definitive endoderm) (**Figure 4a**, **Figure S5**). On Day 0, micropatterned hESCs uniformly expressed SOX2. Following CHIR treatment, TBXT peaked on Day 1, marking the transition to PS cells. SOX17 expression emerged on Day 2 and peaked on Day 3, coinciding with a rapid decline in SOX2 and TBXT (**Figure 4b**). By Days 6–8, expression of these early developmental markers (SOX2, TBXT, SOX17) was minimal. Interestingly, cells in 200 µm micropatterns exhibited a slower decline in SOX2 and TBXT, whereas those in 1000 µm patterns showed a delayed decline in SOX17, suggesting that smaller patterns (200 µm) retain cell pluripotency and delay gastrulation, whereas larger patterns (1000 µm) promote endoderm differentiation.

**Figure 4.**
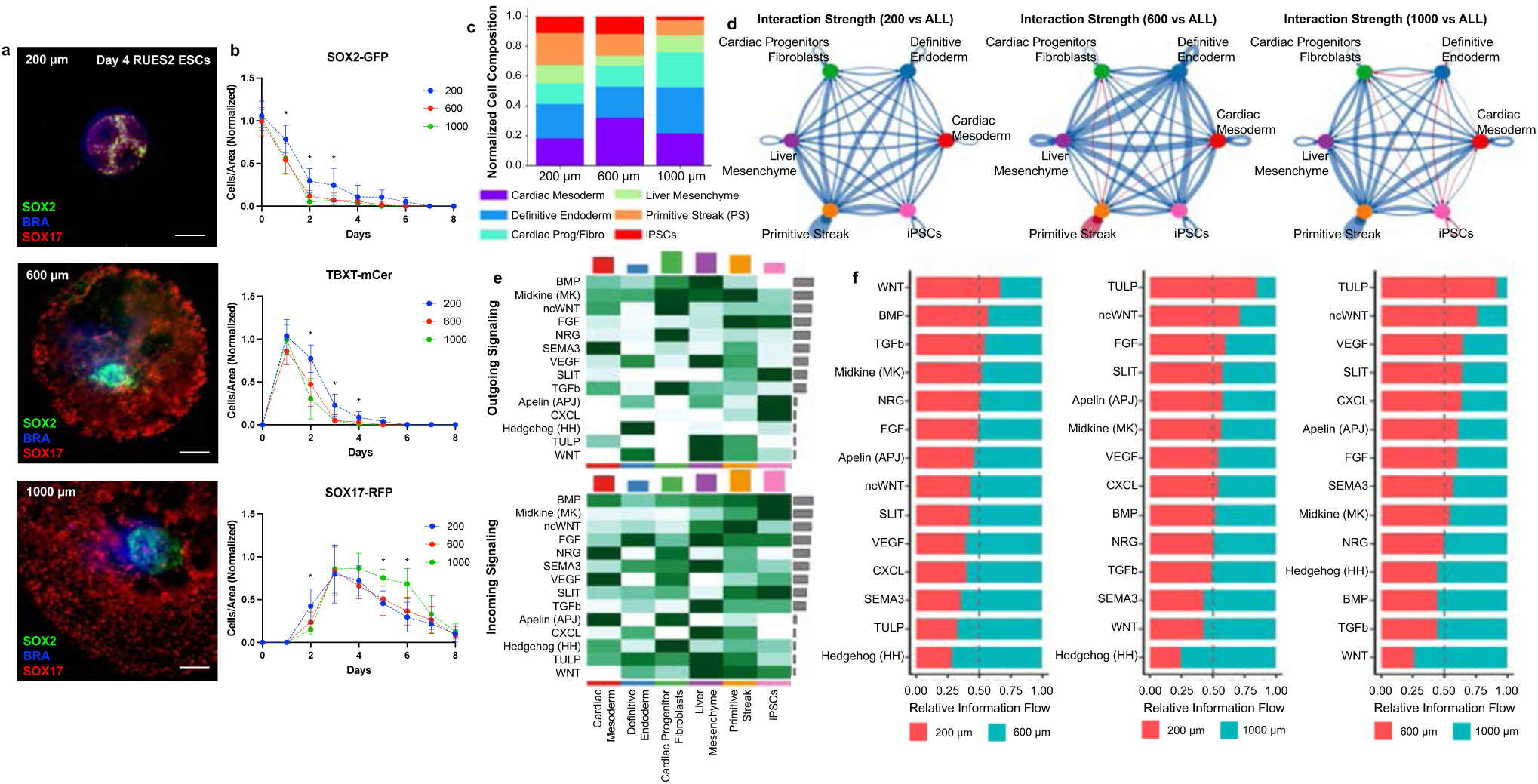
Pattern size modulates lineage dynamics and mesoderm-endoderm crosstalk. (a) Representative images of SOX2–mCitrine, BRA–mCerulean, and SOX17–tdTomato expression in micropatterned RUES2-GLR hESCs on Day 4 of cardioid differentiation. Scale bar: 100 µm. (b) Temporal expression dynamics of SOX2, TBXT (Brachyury), and SOX17 revealed distinct peak expression patterns at different time points (Day 0–Day 8). Sample size > 10 micropatterns from three independent differentiation. Statistical analysis: one-way ANOVA with post-hoc Tukey analysis for each day (*p < 0.05). (c) Comparison of cell composition at Day 4 across three different pattern sizes, illustrating size-dependent lineage specification. (d) Cell-cell interaction strength between six cell types at Day 4, quantified for each pattern size relative to the average interaction strength across all sizes. (e) Outgoing and incoming signaling patterns from six cell types on Day 4, highlighting key signaling pathways involved in early differentiation. (f) Relative signaling strength between two selected pattern sizes for key embryogenesis-associated pathways, demonstrating size-dependent modulation of intercellular communication.

Our scRNA-seq analysis on Day 4 confirmed the trends observed from tri-reporter hESCs: 200 µm patterns retained more iPSCs and PS cells, 600 µm patterns favored cardiac mesoderm, and 1000 µm patterns enriched definitive endoderm **(Figure 4c)**. To understand the cell-cell signaling behind differential mesoderm-endoderm divergence across different pattern sizes, we perform ligand-receptor interaction analysis (CellChat) across six emerging cell types (**Figure 4d**). Increased interaction strength among PS cells, cardiac mesoderm, and cardiac progenitors was observed in 600 µm patterns, suggesting enhanced signaling toward cardiac fate. In contrast, interactions between definitive endoderm and iPSCs were stronger in 1000 µm patterns, suggesting enhanced signaling toward endoderm fate. Of the 23 identified signaling pathways, 10 were selected for further analysis due to their roles in early embryonic development **(Figure S6)**. We found that mesoderm-derived cells were the primary source of non-canonical WNT (ncWNT), NRG, SEMA3, and TGF-β signaling, whereas endoderm-derived cells predominantly contributed to VEGF and Hedgehog (HH) signaling (**Figure 4e**). Furthermore, canonical WNT signaling was significantly inactivated in 600 µm patterns, while ncWNT was highly activated, potentially facilitating transition toward cardiac progenitors (**Figure 4f**). Meanwhile, HH signaling was significantly lower in 200 µm patterns but enriched in 1000 µm patterns, suggesting a crucial role for HH signaling in endoderm induction^43,44^.

### The Impact of Pattern Sizes on Cardioid Cavitation

Previous cardioid models have demonstrated prominent chamber formation^10,13,45–47^. In our study, we observed cavity formation within the cardioids using our deep-tissue imaging technique (**Figure 5a and Figure S7**). Generally, we observed that 200 µm cardioids had limited cavity formation with relatively smaller cavity area (**Figure S7a**), while 1000 µm cardioids had large number of mesh-like interconnected cavities (**Figure S7c**). In contrast, 600 µm cardioids showed large, centralized cavity, resembling of early heart chamber formation (**Figure S7b**). To quantify cardioid cavitation, we measured cavity area, interconnectivity, and number **(Figure 5b-d)**. Smaller (200 µm) cardioids exhibited limited cavity formation compared to 600 µm and 1000 µm cardioids. Although 1000 µm cardioids had more cavities, their average cavity size appeared smaller than that of 600 µm cardioids.

**Figure 5.**
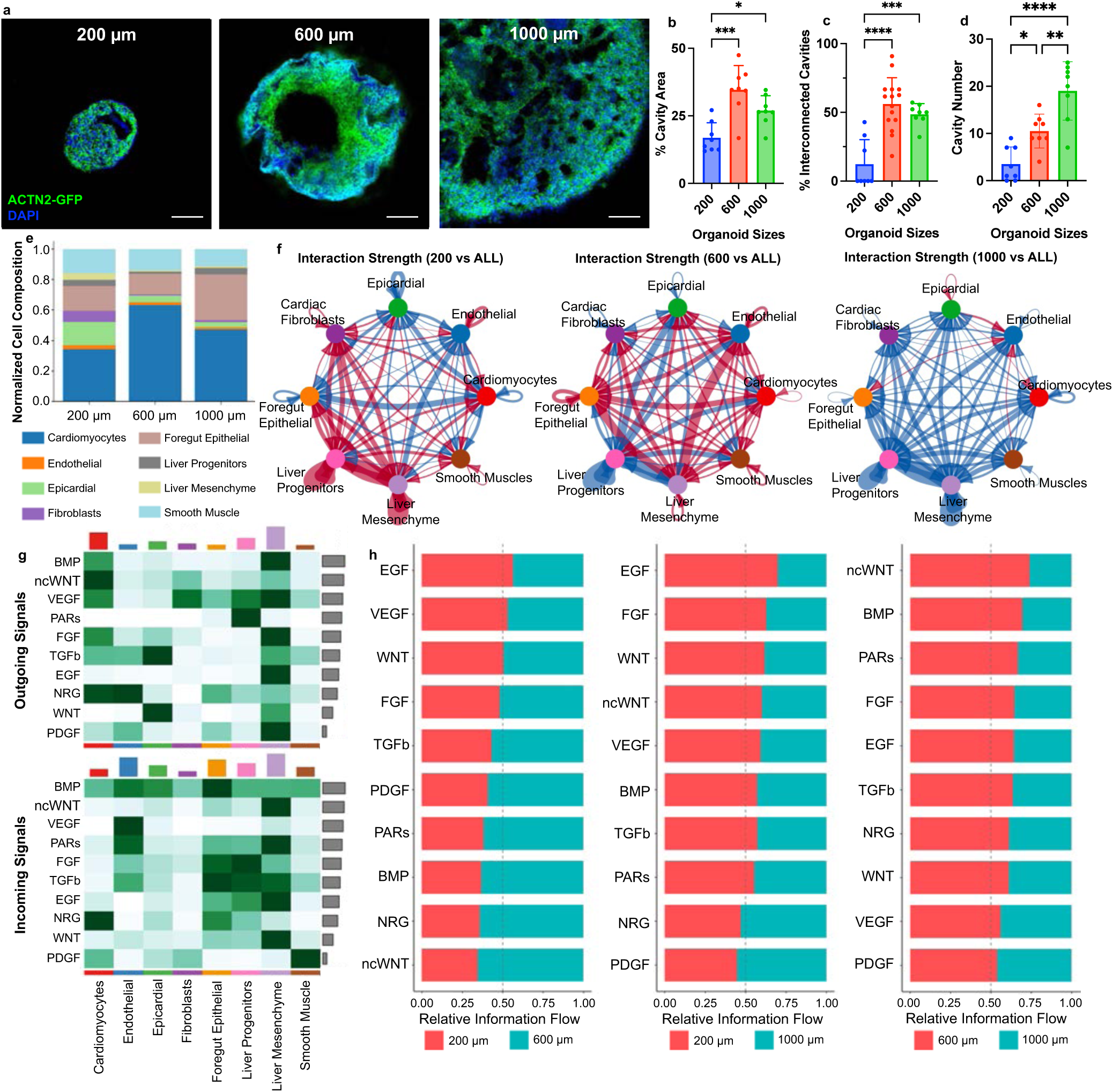
Pattern sizes modulate cardioid cavitation and heart-foregut crosstalk. (a) Representative images of internal cavitation of Day 21 cardioids generated from different pattern sizes. Scale bar: 100 µm. Quantification of cardioid cavitation based on (b) cavity area relative to cardioid size, (c) interconnected cavities relative to total cavities, and (d) total cavity numbers showed enhanced cavitation in 600 µm cardioids. Sample size > 6 cardioids from three independent differentiation. Statistical analysis: one-way ANOVA with post-hoc Tukey analysis (*p < 0.05, **p<0.01, ***p<0.001, ****p<0.001). (e) Comparison of cell composition at Day 21 across three different cardioid sizes. (f) Cell-cell interaction strength between eight cell types at Day 21, quantified for each pattern size relative to the average interaction strength across all sizes. (g) Outgoing and incoming signaling patterns from eight cell types on Day 21, highlighting key signaling pathways involved in heart-foregut co-development. (h) Relative signaling strength between two selected pattern sizes for key organogenesis-associated pathways, demonstrating size-dependent modulation of heart-foregut crosstalk.

To assess contractile function, we generated cardioids from a GCaMP6f-reporter hiPSC line and stained them with a membrane potential dye (PhoS VSD-Cy5). Brightfield (**Figure S8a, Movie S3**), GCaMP6f-GFP (**Figure S8b, Movie S4**), and VSD-Cy5 (**Figure S8c, Movie S5**) recordings from the same cardioids were used for contractile motion, calcium transient, and action potential analyses. From motion analysis, we found no significant difference on beat rate, contraction velocity, and relaxation velocity across pattern sizes, but 1000 µm cardioids showed prolongated time interval between contraction and relaxation. The elongation of contraction duration in 1000 µm cardioids also reflected as longer calcium decay time (t30, t50, t75) in calcium transient analysis and longer action potential duration (rising time, APD30, APD50, APD80) in membrane potential analysis. Notably, 600 µm cardioids showed the shortest calcium decay times (t50 and t75), indicating an enhanced calcium handling ability. These structural and functional analyses suggest that micropattern size modulates cardioid cavitation and contractile properties, with 600 µm cardioids exhibiting prominent chamber formation and enhanced contractile function.

Our scRNA-seq analysis on Day 21 revealed that 600 µm cardioids had the highest cardiomyocyte population (∼65%) compared to 200 µm (∼35%) and 1000 µm (∼50%) cardioids (**Figure 5e**). 200 µm cardioids were enriched in cardiac stromal cells (epicardial and smooth muscle cells), while 1000 µm cardioids contained a higher proportion of endoderm cells. To understand the cell-cell signaling behind enhanced cavitation and function in 600 µm cardioids, we performed CellChat analysis (**Figure S9**). 600 µm cardioids were found to have stronger interactions between foregut epithelial cells and cardiomyocytes (**Figure 5f).** Endoderm cells were the primary outgoing signal sources for VEGF and PARs, both of which are key regulators for angiogenesis of endothelial cells (**Figure 5g**). Notably, liver mesenchymal cells emerged as a central signaling hub, mediating interactions between heart and liver cells within the cardioids. Additionally, key cardiogenic pathways such as NRG and ncWNT were enriched in 600 µm cardioids, comparing to 200 cardioids and 1000 µm cardioids (**Figure 5h**).

## DISCUSSION

### Mesoderm-Endoderm Reciprocal Signaling in Heart Development

During heart development, the mesoderm and endoderm engage in reciprocal signaling interactions that drive lineage specification and organogenesis, as demonstrated across multiple species^48^. The mesoderm provides essential signals that guide endodermal fate, instructing foregut endoderm to develop into hepatic tissues while also contributing to the development of connective tissue and vascular progenitors in endoderm-derived organs^49,50^. FGF signaling from the cardiac mesoderm has been shown to facilitate liver formation, while in return, hedgehog (HH) signaling from the ventral foregut endoderm plays a crucial role in modulating splanchnic mesoderm patterning for proper heart morphogenesis^51,52^. In a multi-lineage organoid model, co-developing cardiac and intestinal tissues gradually segregated into distinct compartments at opposite ends of the organoid, separated by intervening mesenchymal tissues. This mesoderm-endoderm co-development promoted the maturation of both cardiac and intestinal tissues^17^. Compared to our cardioid model with foregut development, the multi-lineage organoid model exhibited prominent midgut differentiation, likely due to the addition of high-dose ascorbic acid.

From our deep-tissue imaging results, we observed the present of endoderm cells inside the cardioids with two main phenotypic structures: cavity-lining epithelium that represent the development of foregut lumen, while rosette-like endoderm islands that indicate the early development of liver bud. The co-emergence of mesoderm-heart and endoderm-liver lineages was further confirmed by our scRNA-seq results. Transcriptional profiles of paracrine signaling further elucidate reciprocal mesoderm-endoderm interactions at Day 4 and the coordinated signaling for heart-liver co-development at Day 21. Endoderm differentiation was highly dependent on FGF signaling, initially provided by primitive streak (PS) cells at Day 4 and later shifting to cardiomyocytes and liver mesenchymal cells at Day 21. Definitive endoderm was the primary signaling source of HH, which was critical for cardiac progenitor specification at Day 4. This aligns with previous single-cell transcriptomic analyses highlighting the essential role of endoderm-derived HH signaling in the development of cardio-pulmonary progenitors at the interface of the second heart field and the ventral foregut region^53^. Furthermore, foregut epithelial cells at Day 21 are essential sources of VEGF, NRG, and PAR signaling, which are critical for myocardial cavitation.

### Size modulation of embryonic development

Controlling the differentiation of PSCs through micropatterning has emerged as a powerful in vitro approach for studying stem cell fate decisions, mammalian embryogenesis, and spatial patterning^54,55^. In a micropatterned human gastruloid model with spatially organized germ layers, reducing pattern size led to the loss of SOX2-expressing ectodermal populations at the colony center^56^, shifting from ectodermal cells to mesendodermal derivatives^57^. In a micropatterned spinal cord organoid model, BMP and HH signaling was modulated by pattern sizes that resulted in the change in spatial patterning of dorsoventral axis^58^. Our previous study has reported that cell collective behaviors within micropatterned hiPSC colonies played a critical role in directing the spatial differentiation during mesoderm induction^24^. Here, we observed smaller 200 µm patterns had higher SOX2 expression and larger 1000 µm patterns had higher SOX17 expression at Day 4. Furthermore, pattern sizes had a strong impact on cavitation of cardioids and heart-foregut interactions at Day 21. 600 µm cardioids showed favorable cardiomyocyte differentiation and prominent chamber formation. More importantly, 600 µm cardioids showed a clear enhancement in BMP, NRG, and ncWNT signaling comparing to 200 µm and 1000 µm cardioids, and these signaling pathways are critical for myocardial cavitation and compaction.

### Myocardial Cavitation in Cardioids

During heart chamber development, cardiac tissue undergoes a complex morphogenetic process to establish the distinct chambered structure with compacted myocardial tissues. The self-organizing nature of cardioid offers great potential in modeling heart chamber formation and understanding the signaling mechanisms governing cavitation and chamber morphogenesis^16,17^. Our micropatterned cardioids showed a direct relationship between initial tissue dimensions and subsequent cavitation levels. Specifically, small (200 µm) cardioids exhibited single but tiny cavities, large (1000 µm) cardioids developed a greater number of small, interconnected cavities, whereas mediate (600 µm) cardioids developed larger, singular cavities resembling early heart chamber formation. Our previous publications demonstrated that 600 µm pattern size results in higher volume of cardioids^59^, allowing to develop larger cavity spaces. In contrast, 1000 µm pattern size intent to lead to flatten tissue morphology similar to 2D cardiac differentiation, leading to a mesh-like spongy network inside the cardioids. These results indicated that cavitation process might be highly associated with the geometric volume of the cardioids. There might be a volumetric threshold that switch from singular chamber development to mesh-like cavitation, which is worthy to further evaluated in the future studies.

Our findings reveal that cavitation in micropatterned cardioids is strongly influenced by initial tissue dimensions, highlighting a geometric control over morphogenetic outcomes. Small (200 µm) cardioids produced only tiny single cavities, whereas large (1000 µm) cardioids exhibited multiple small, interconnected cavities with a spongy, mesh-like morphology, since 1000 µm pattern size intent to lead to flatten tissue morphology similar to 2D cardiac differentiation. By contrast, intermediate-sized (600 µm) cardioids generated larger, singular cavities resembling early heart chamber formation, consistent with our previous observations that this pattern size develops greater tissue volume. These results suggest that cavity formation may be governed by a volumetric threshold, where sufficient tissue mass promotes the emergence of singular chamber-like structures, while exceeding this threshold drives a transition toward disorganized, mesh-like cavitation. This volumetric dependence underscores the importance of geometric constraints in shaping self-organizing cardiac morphogenesis and raises the possibility that manipulating tissue dimensions could be a powerful strategy to steer chamber formation in vitro. Future studies will be needed to test the existence of such thresholds and to define how tissue geometry integrates with molecular signaling pathways to guide cavity morphogenesis.

From our scRNA-seq analysis, NRG signaling has been hypothesized as a key mediator of crosstalk between cardiomyocytes and endothelial cells. NRG signaling is a well-established regulator of myocardial cavitation and compaction, essential for the transition from a primitive, spongy myocardium to a more structured, mature heart wall^60–63^. Foregut epithelial cells not only lined the cavities within cardioids, but they also acted as a secondary signaling source for VEGF and NRG, potentially serving as a decoy role for limited population of endothelial cells within our cardioids. This further underscores the importance of multi-lineage interactions in cardioid morphogenesis, reinforcing the hypothesis that cardiac chamber formation is not solely an intrinsic myocardial process but is heavily shaped by surrounding mesodermal and endodermal tissues.

### Limitations of the study

One key limitation of this study is that our cardioid differentiation protocol relies on WNT signaling modulation using two small molecules. Other protocols employing complex growth factor combinations have successfully generated cardioids with single large chambers^13^. However, these methods appear to limit epicardial and endodermal differentiation, thus constraining our ability to investigate heart-foregut developmental interactions. Our simplified differentiation protocol, on the other hand, highlights how biophysical cues influence on the cardioid cavitation and chamber formation. A previous study also reported endocardial tissue (NFATC1+) as a distinct layer independent of the endothelial vascular network (PECAM1+)^10^. While NFATC1 is highly expressed in endocardial cushions during valve formation^64,65^, its expression is minimal in the endocardial cells from heart’s free walls^34^. However, we did not detect NFATC1+ endocardial cells in our scRNA-seq data. Future studies should explore strategies to enrich NFATC1+ endocardial cells in our cardioids, enabling a more comprehensive model of early heart development.

Another challenge in cardioid models is the variability in cellular composition and maturation. Since cardioids rely on self-organization with minimal external guidance, they often exhibit heterogeneity across different batches and different iPSC lines. This variability can arise from differences in intrinsic genetic programs, epigenetic states, and responsiveness to signaling cues. For example, previous research has shown that different iPSC lines exhibited different responses to IGF1 and BMP4 activation during retinal organoid differentiation^66^, highlighting the need to account for cell-line specific variability in organoid models. In this study, cardioids were generated from multiple hiPSC lines carrying different reporter systems; however, these cell lines share the same WTC genetic background. While our previous work successfully derived cardioids from a different hiPSC line^59^, future studies should incorporate a broader panel of hiPSC lines from healthy donors with diverse genetic backgrounds, including variations in sex, ethnicity, and ancestry. Addressing batch-to-batch variability will require optimizing culture conditions, standardizing differentiation protocols, and integrating engineering controls and computational models for identifying and mitigating sources of technical variation.

## METHODS

### Micropatterning of tissue culture surfaces

Surface micropatterning on tissue culture polystyrene was performed using a selective etching approach as previously described^25^. Patterned wafers (SU8 masters) were fabricated using standard SU8 photolithography to create molds with raised pattern features. Poly(dimethyl siloxane) (PDMS) was prepared at a 10:1 wt/wt ratio of elastomer base to curing agent, cast onto the SU8 masters, and clamped using clear transparency sheets and glass slides. This process produced thin PDMS stencils with through-holes corresponding to the raised patterns on the SU8 master molds. A non-fouling poly(ethylene glycol) (PEG) solution was prepared by mixing 150 mg PEG 1000 (Polysciences), 1.8 mL PEGDA 400 (Polysciences), 14.55 mL isopropyl alcohol, and 0.45 mL Milli-Q water. The solution was applied to 6-well tissue culture plates and cured under UV light exposure (Dymax UV Illuminator; model no. 2000EC) for 45 seconds. Micropatterns were then generated via selective oxygen plasma etching (Oxygen plasma treatment system, PlasmaEtch PE50XL) of the PEG coating using the PDMS stencils as masks. Following fabrication, the micropatterned tissue culture plates were sterilized by immersion in 70% ethanol for 1 hour, followed by thorough washing with sterile phosphate-buffered saline (PBS) to ensure biocompatibility before cell culture experiments.

### Cell Lines

The wild-type (WTC) hiPSC line (RRID: CVCL_0600) was obtained from Dr. Conklin’s laboratory at the Gladstone Institute of Cardiovascular Research. This hiPSC line was originally derived from a skin biopsy of a healthy adult Asian male donor in his early thirties. Reprogramming of the original fibroblasts was performed using episomal methods with the transcription factors LIN28A, MYC (c-MYC), POU5F1 (OCT4), and SOX2. The hiPSC reporter lines used in this study were engineered by the Allen Institute using the WTC hiPSC line (RRID: CVCL_0600) as the parental cell line and were purchased from the Coriell Institute. These reporter lines are fluorescently tagged with mEGFP targeting ACTN2 (sarcomere alpha-actinin) and TNNI2 (troponin I), enabling visualization of cardiomyocyte structures. Additionally, the RUES2-GLR hESC line was obtained from Dr. Lian’s laboratory as part of this collaborative effort to evaluate early germ cell specification. These cells were engineered from RUES2 hESC line (RRID: CVCL_5323) using CRISPR-Cas9 technology with three fate reporters: SOX2– mCitrine, BRA–mCerulean, and SOX17–tdTomato^67^. All cell lines used in this study are contamination-free. The parental lines (WTC and RUES2) are widely recognized and consistently employed across the stem cell field to ensure reliable results. From these well-characterized parental lines, we generated fluorescent reporter lines that are uniquely suited for deep-tissue and live-cell imaging of cardioids.

### hPSC maintenance

Pluripotent stem cells (PSCs), including hiPSCs and hESCs, were cultured following standard PSC maintenance protocols. Cells were maintained in Essential 8 (E8) medium (Thermo Fisher, cat. no. A1517001) and passaged at a 1:6 ratio approximately every three days upon reaching at least 80% confluency. For passaging, cells were dissociated using Accutase (Thermo Fisher, cat. no. A1110501) for 3–4 minutes and then neutralized with E8 medium. The cell suspension was centrifuged, and the resulting pellet was resuspended in fresh E8 medium for replating.

### Generation of cardioids

Micropatterned surfaces were coated with diluted Geltrex hESC-qualified matrix (Thermo Fisher, cat. no. A1413302) and incubated at 37°C for 1 hour before cell seeding. hiPSCs were maintained using standard PSC culture practices in E8 medium. PSCs were seeded at a density of 6.0 × 10⁵ cells per well in a micropatterned 6-well plate (∼0.63 × 10⁵ cells per cm²), supplemented with 10 µM Y27632 (Stem Cell Technologies, cat. no. 72304). Cardiac differentiation was initiated when the micropatterns reached confluency, approximately three days after seeding, and was induced via small molecule modulation of the Wnt/β-catenin pathway^68^. Differentiation was initiated with the GSK3 inhibitor CHIR99021 (12 µM) on Day 0 (Stem Cell Technologies, cat. no. 72054) and followed by the WNT pathway inhibitor IWP4 (6 µM) on Day 2 (Stem Cell Technologies, cat. no. 72552). Small molecules were diluted in RPMI 1640 medium (Thermo Fisher, cat. no. 11875093) supplemented with B27-minus insulin (RPMI/B27-minus insulin) (Thermo Fisher, cat. no. A1895601). Cardioids began spontaneous contraction around Day 9 of differentiation and were subsequently maintained in RPMI 1640 medium supplemented with complete B27 supplement (RPMI/B27 Complete) (Thermo Fisher, cat. no. 17504044) until Day 21.

### Immunostaining and tissue clearance

Cardioids were characterized using immunofluorescence staining for cardiac and endodermal markers (Table S1). Samples were sacrificed and fixed with 4% (vol/vol) paraformaldehyde (PFA) for 10 minutes, followed by washing and permeabilization with 0.2% (vol/vol) Triton X-100. Blocking was performed using 2% (wt/vol) bovine serum albumin (BSA), and samples were incubated with primary antibodies at the appropriate dilution for 1 hour at room temperature. After incubation, the primary antibody was removed, and samples were washed with PBS before incubation with secondary fluorescent antibodies in the dark for 2 hours. Nuclei were stained with 300 nM DAPI.

To enhance deep-tissue imaging, tissue clearing was performed using the CUBIC method^69^. Cardioids were incubated in Reagent 1 for three days to remove lipids, facilitating tissue transparency. Subsequently, cardioids were transferred to Reagent 2, which adjusts the tissue refractive index to match the imaging medium, allowing for high-resolution imaging in combination with immunofluorescence staining and reporter cell line visualization. <u>Reagent 1:</u> 25 wt% urea, 25 wt% N,N,N’,N’-tetrakis(2-hydroxypropyl)ethylenediamine, 15 wt% Triton X-100, and 35 wt% PBS. <u>Reagent 2:</u> 50 wt% sucrose, 25 wt% urea, 10 wt% 2,2’,2’’-nitrilotriethanol, 0.1% (vol/vol) Triton X-100, and 15 wt% PBS.

A Zeiss LSM U880 confocal microscope, equipped with a multi-photon laser source (SpectraPhysics MaiTai laser with 700–1000 nm multiphoton excitation), was used for both confocal and two-photon imaging. Imaging was performed using a 20× water-dipping objective (W Plan-Apochromat, Zeiss) to capture z-stacks with an 8 µm spacing between slices. These z-stacks were used for 3D reconstruction and visualization of internal cardioid structures.

### Image analysis for tri-reporters and cardioid cavitation

The RUES2-GLR cell line was utilized to quantify early germ cell development. Daily images were acquired over an 8-day period and processed in ImageJ, where noise removal, thresholding, and watershed segmentation were applied to isolate and identify individual cells. Cell counting was performed using the Analyze Particles function in ImageJ, and the counts were normalized to the micropattern area.

Two-photon imaging was employed to quantify the presence of cell-void cavities within the cardioids, following previously published methods^10^. Individual slices from the z-stack were converted to binary images in ImageJ. Cavities were quantified by measuring the percentage of white space relative to the overall tissue area. The tissue region was first delineated using the Polygon Selections tool, and its area was determined using the Measure function under Analyze. Cavitation was quantified as the percentage of the non-tissue (white) area within the outlined cardioid region

### Function Analysis for Cardioids

Contraction physiology was assessed by evaluating contraction motion, calcium transient, and action potential. GCaMP6f hiPSC-derived cardioids were imaged in a controlled environment using an on-stage microscope incubator (OkoLab Stage Top Incubator, UNO-T-H-CO2) set to 37°C and 5% CO₂ to maintain physiological conditions. Imaging was conducted on a Nikon Ti-E inverted microscope equipped with an Andor Zyla 4.2+ digital CMOS camera. Videos of contracting cardiac organoids were captured in brightfield at 60 frames per second for 10 seconds, then exported as a series of single-frame image files for contractile motion analysis using an open-source MATLAB motion tracking algorithm^70^. Calcium flux videos were captured under 488 nm excitation at 33 frames per second. Imaging analysis was conducted using an open-source MATLAB algorithm, which detected calcium flux signals from fluorescence traces associated with manually drawn regions of interest (ROIs)^71^. Fluorescence bleaching decay was corrected, and key parameters were calculated and corrected based on the beat rate. The upstroke duration (UPD) represents the time it takes for the calcium flux to reach peak fluorescence intensity, while t30, t50 and t75 indicate the time it takes for the flux to decay by 30%, 50% and 75% of the peak value, respectively. For action potential analysis, beating cardioids were stained with VoltageFluor/BeRST membrane potential dye (1:100) for 20 minutes, and then washed twice with PBS and refreshed with new RPMI/B27 complete media. Videos were recorded under 670 nm excitation at 33 frames per second. Action potential signals were analyzed using an open-source MATLAB algorithm^71^. Action potential repolarization was determined by the fluorescent signal, where rising time represents the time to peak fluorescence intensity, and APD30, APD50, APD80 correspond to the time duration for action potential duration at 30%, 50%, and 80% of peak value, respectively.

### Statistics

All data are presented as mean ± standard deviation. At least three biological replicates were included in each group from independent differentiation. At least 3 cardioids were analyzed for each differentiation batch as technical replicates. Statistics are analyzed using one-way ANOVA with Tukey post-hoc test. All analysis was performed using Prism 10 (GraphPad). Values of *p* < 0.05 were considered statistically significant.

### Single-cell RNA-sequencing, alignment, quality control, and preprocessing

Cardioids were isolated by first scraping, aspirating, and discarding residual cells and tissue within the wells. Only beating cardioids (∼ 15 – 25 cardioids) were collected from one well of 6-well plates. For each sample, at least 2 wells were used to pool micropatterned cells or cardioids for scRNA-seq experiments. For timepoints prior to Day 6, samples were dissociated with Accutase and quenched with RPMI/B27 minus insulin medium, while samples from Day 6 onward were dissociated using cardiomyocyte dissociation medium (Stem Cell Technologies, cat. no. 05025) and quenched with RPMI/B27 complete medium. Cells were singularized by repeated pipetting and passing through a cell strainer. Single-cell RNA sequencing libraries were prepared using the Chromium Single Cell 3′ reagent kit v3 (10x Genomics, PN-1000075) according to the manufacturer’s protocol. Cells were diluted into the Chromium Single Cell A Chip to obtain approximately 6,000 single-cell transcriptomes. After preparation, libraries were sequenced on a NextSeq 500 (Illumina) with 75-cycle high-output kits (Index 1 = 8, Read 1 = 26, and Read 2 = 58).

Sequencing data were trimmed and quality-filtered using cutadapt (v4.1). Read 2 was trimmed by removing poly(A) and poly(G) sequences from the three-prime end, and removing template switch oligonucleotide sequence and its reverse complement from the five-prime end. After trimming, all reads shorter than 18bp were discarded. The remaining data were aligned and quantified using STARsolo (STAR v2.7.10a). The filtered count matrices (-- soloCellFilter EmptyDrops_CR, --soloFeatures GeneFull) were used for downstream analysis. For preprocessing of all samples, SoupX (v1.5.2) with default settings was used to remove ambient RNA contamination^72^. Further quality control and preprocessing were performed individually for each sample using Scanpy (v1.9.1)^73^. First, cells with fewer than 500 detected features or fewer than 1,500 unique molecules were excluded. Genes detected in fewer than five cells were also removed. Cells with more than 20% of unique molecules mapping to the mitochondrial genome were also excluded. Doublet identification was performed using Scrublet (v0.2.3)^74^. Cells with a predicted doublet score greater than 0.4 were removed from Day 0 and Day 1 samples, while cells with a predicted doublet score greater than 0.2 were removed from all other samples. After filtering and doublet removal, a total of 96,401 cells were retained for subsequent analysis (**Figure S3a**).

### Single-cell RNA-sequencing, trajectory construction, pseudo-time analysis, and cell-cell interaction analysis

After quality control, the samples were merged. Gene expression data was normalized, log-transformed, and scaled. The top 2,000 most variable genes were identified and used for principal component analysis (PCA). The cell cycle phases were inferred using the cell cycle function from Scanpy. To remove batch effects and cell cycle differences, batch correction was performed on the first 50 principal components using Harmony (harmonypy, v0.0.6)^26^, with both sample and cell cycle phase included as covariates (**Figure S3b**). The dimensions of the Harmony embedding were reordered by decreasing variance, after which a nearest neighbor graph was computed and Leiden clustering was performed with a resolution of 1.0. The resulting clusters were labeled and merged based on a set of curated canonical gene markers. The dimensions containing 95% of the total variance within the Harmony embedding were reduced using PHATE (Potential of Heat-diffusion for Affinity-based Transition Embedding) (v1.0.9) to build the lineage trajectory of heart-foregut co-development^27^. Partition-based Graph Abstraction (PAGA) generalized the relationships between cells clusters to quantify the connectivity between each cluster^75^. Force-directed graphs were built in ScanPy utilizing a force atlas layout to construct the diffusion pseudo-time, with the root clusters that represent the possible position of cell differentiation along its developmental trajectory compared to all cells within the graph^76,77^. A deeper investigation of single-cell fate of cardiomyocyte maturation was conducted with CellRank^33^. Cell-cell crosstalk was conducted in R utilizing the CellChat library, which utilizes a human ligand-receptor interaction database to calculate the biologically significant cell-cell communication network^78^.

## SUPPLEMENTARY INFORMATION

Supplementary information can be found online.

## ACKNOWLEDGEMENTS

This work was supported by the NIH [R01HD101130, R21HD11458, R15HD108720, R01HL175050, R56DK133147], the NSF [CMMI-2130192 and CBET-1943798], and AHA [24IVPHA1288417].

## AUTHOR CONTRIBUTIONS

P.H., A.K. and Z.M. conceived the study. D.W.K. and A.K. performed the computational work for single-cell RNA sequencing data and bioinformatics analysis. P.H., N.Y.M. and M.C. performed all experiments, collected raw data, and processed the recorded videos. X.L.L shared the RUES2-GLR human ESC line and instructed the use of this line for tri-lineage differentiation analysis. Y.Z., N.T., and I.D.V. provided insights on single cell transcriptomics on early embryogenesis and heart development. J.A. and H.Y. provided insights on cardioid development and physiology, B.D.C. and Z.M. supervised the project development. P.H. D.W.K., A.K., N.Y.M. and Z.M. wrote the manuscript with discussion and improvements from all authors. Z.M. funded the study.

## RESOURCE AVAILABILITY

The code for preprocessing and analysis of the single-cell count matrices is available at https://github.com/mckellardw/scCardiacOrganoid.

The raw single-cell RNAseq data is available through NIH GEO (GEO accession ID is GSE293554) https://www.ncbi.nlm.nih.gov/geo/query/acc.cgi?acc=GSE293554

Further information and requests for resources and reagents should be directed to and will be fulfilled by the lead contact, Zhen Ma.

## SUPPLEMENTARY INFORMATION

**Figure S1.**
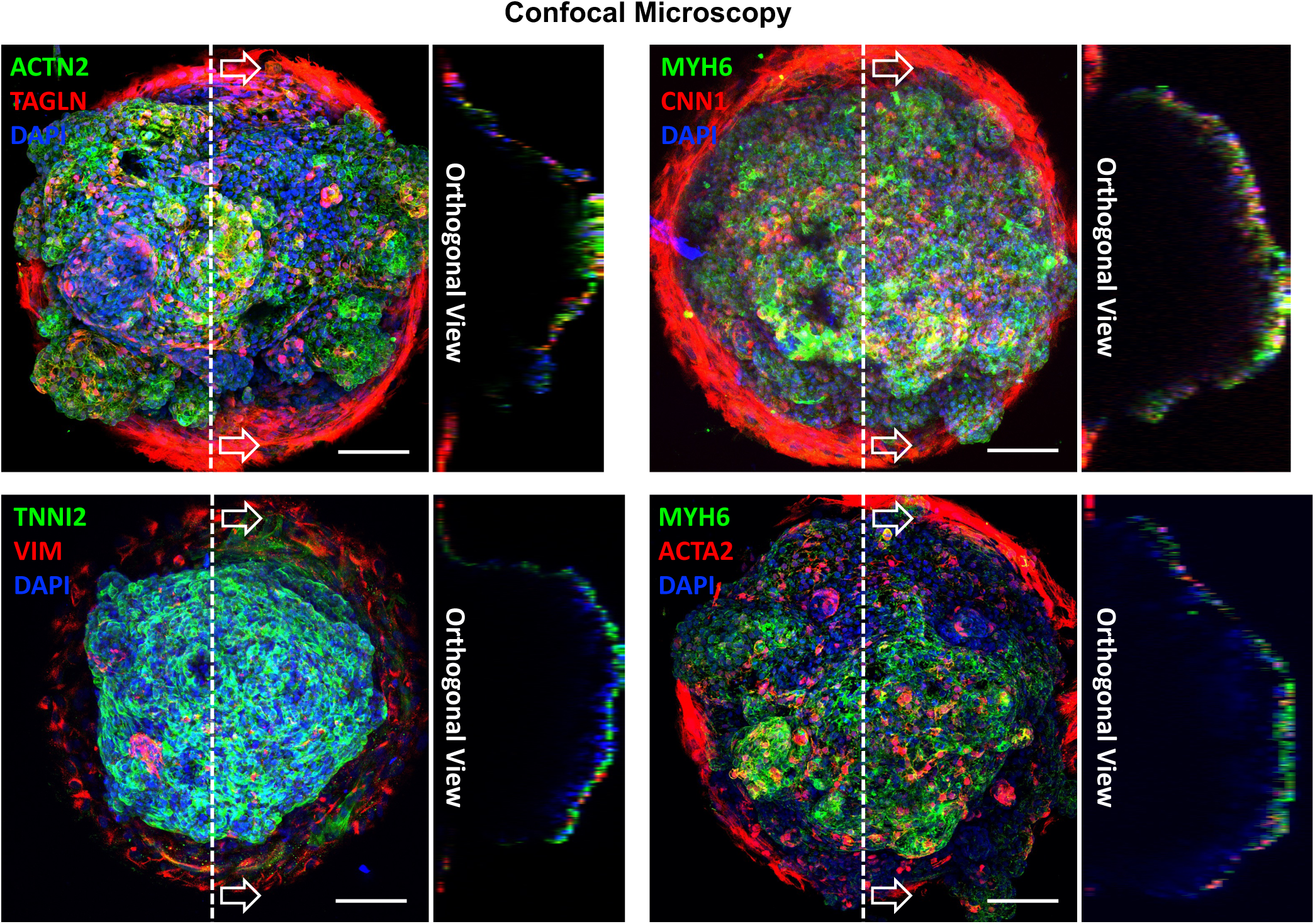
Representative 3D reconstruction images of confocal microscopy of micropatterned cardioids (related to Figure 1). The micropatterned cardioids were derived from WTC hiPSC line without any fluorescent reporter, whole mounted without cryo-sectioning, and stained with cardiomyocyte markers (TNNI2, ACTN2, and MYH6) and stromal cell markers (TAGLN, CNN1, VIM, and ACTA2). Scale: 100 µm.

**Figure S2.**
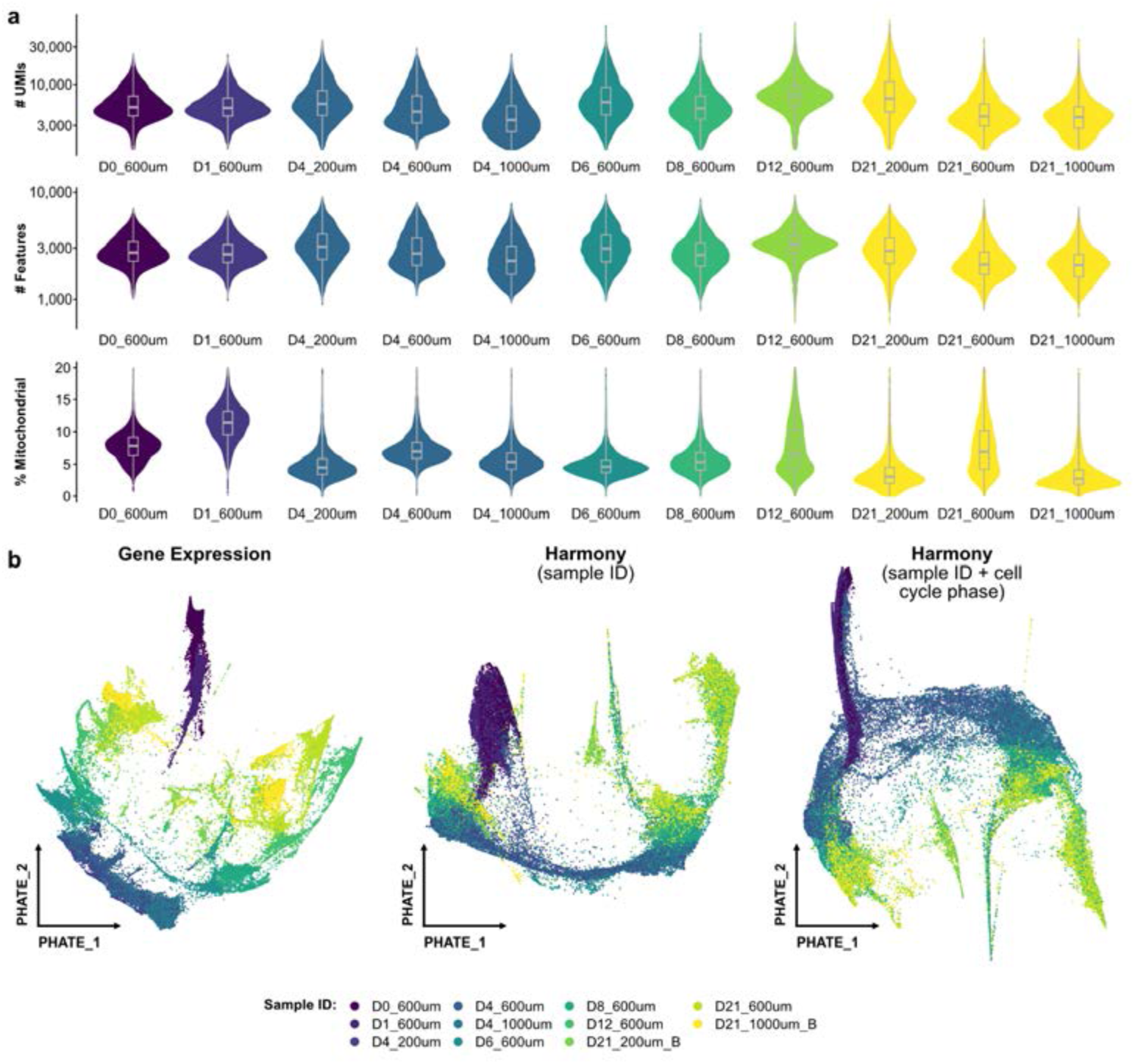
Quality control of scRNA-seq dataset (Related to Figure 2). (a) The number of unique molecular identifier (UMI), number of features, and % of mitochondrial genes were analyzed as quality control metrics in preprocessing steps. (b) Comparison of trajectory construction with no batch correction, batch correction based on sample ID, and batch correction based on sample ID and cell cycle phases.

**Figure S3.**
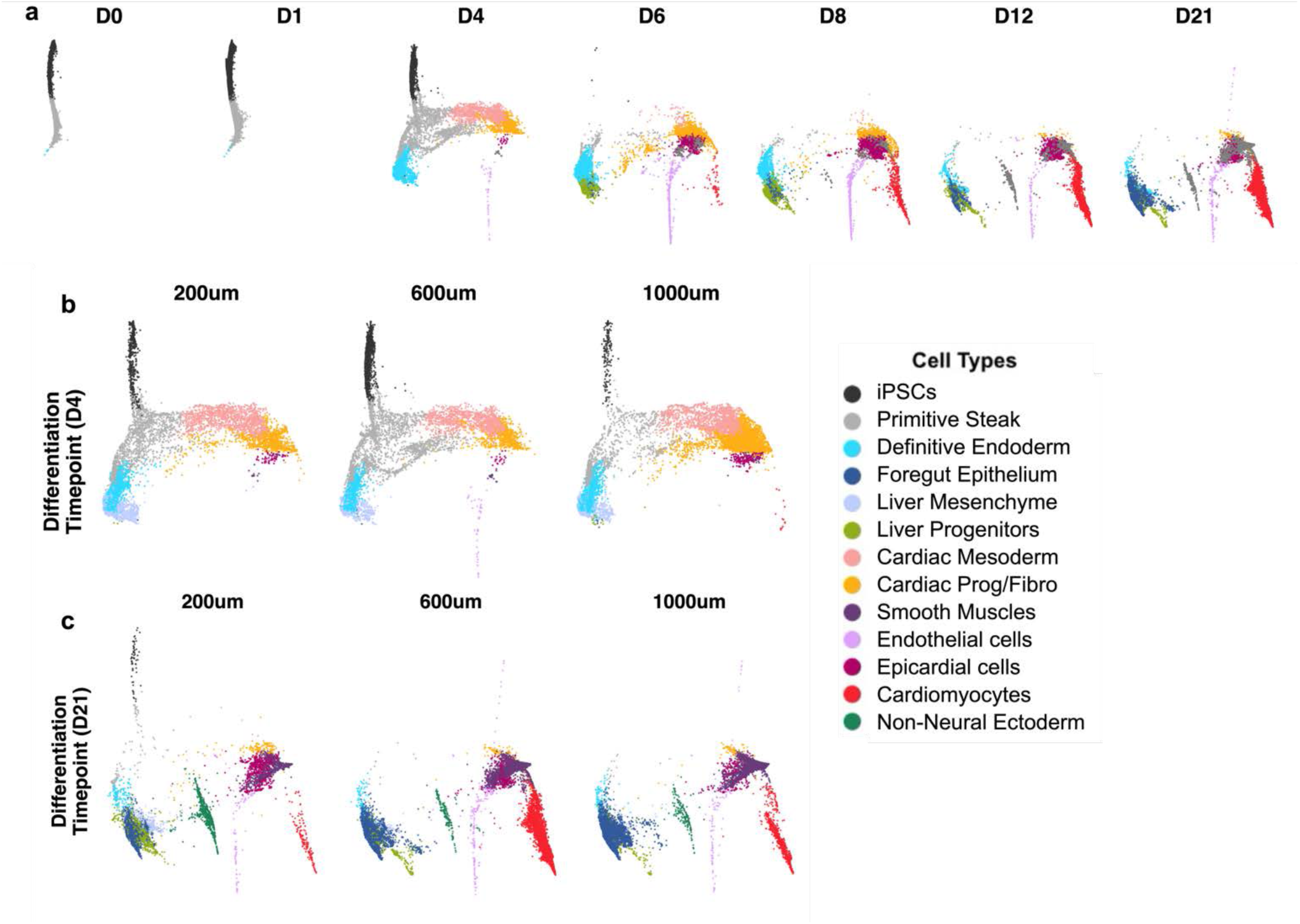
PHATE Embedding for Individual Samples (Related to Figure 2). (a) PHATE embedding depicting developmental progression at seven time points during 600 µm cardioid differentiation. (b) PHATE embedding at Day 4, illustrating mesoderm-endoderm divergence from three different pattern sizes. (c) PHATE embedding at Day 21, visualizing heart-foregut co-development in cardioids derived from three different pattern sizes.

**Figure S4.**
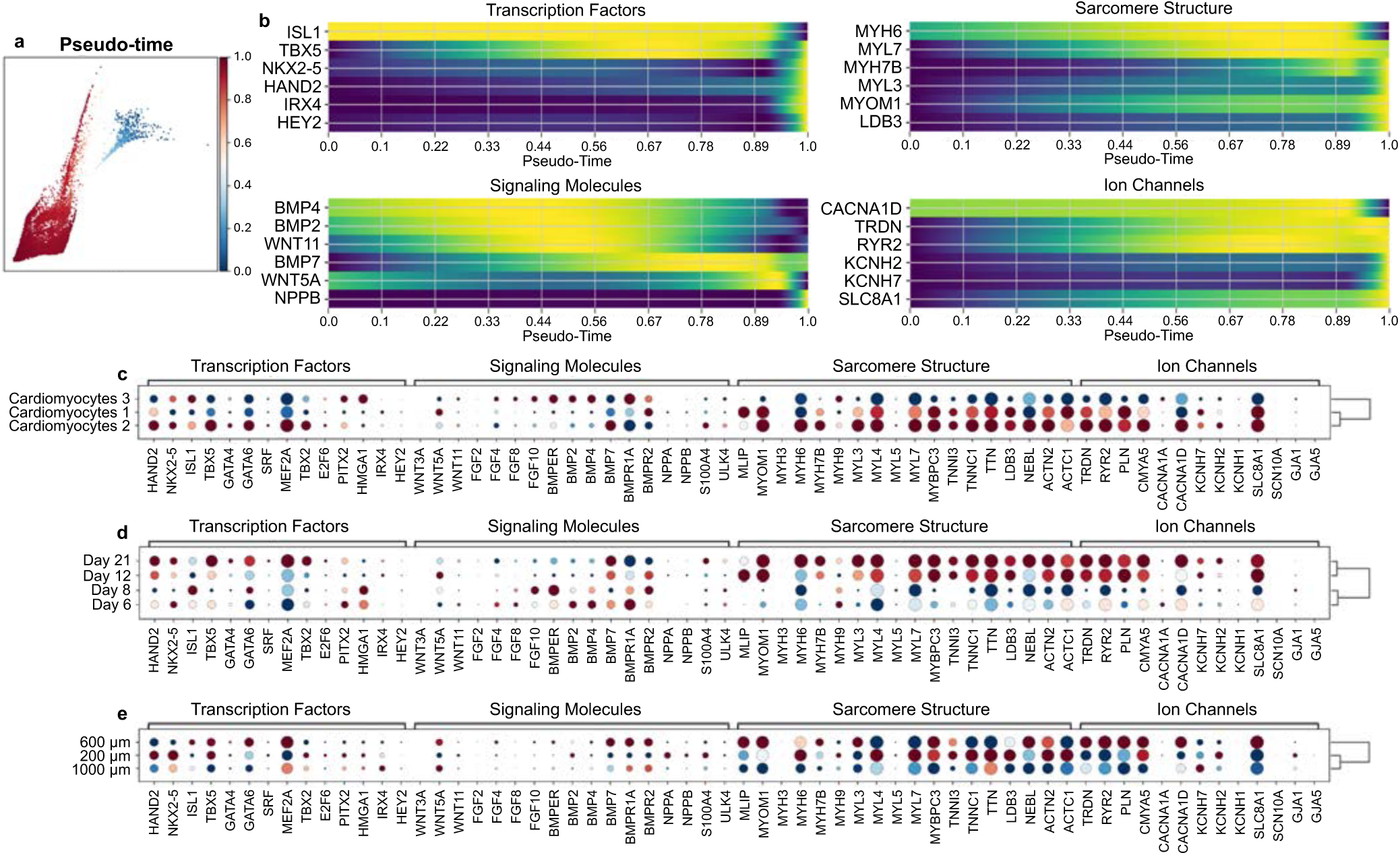
Profile cardiomyocyte differentiation progress (Related to Figure 3). (a) Pseudo-time was constructed for cardiomyocyte clusters using CellRank. (b) Pseudo-time progression of key cardiomyocyte genes for transcription factors, signaling molecules, sarcomere structures, and ion channels. Gene expression comparison for (c) three cardiomyocyte clusters from PAGA-guided force-directed dimensions, (d) four differentiation time points during cardioid differentiation, and (e) Day 21 cardioids generated from three different pattern sizes.

**Figure S5.**
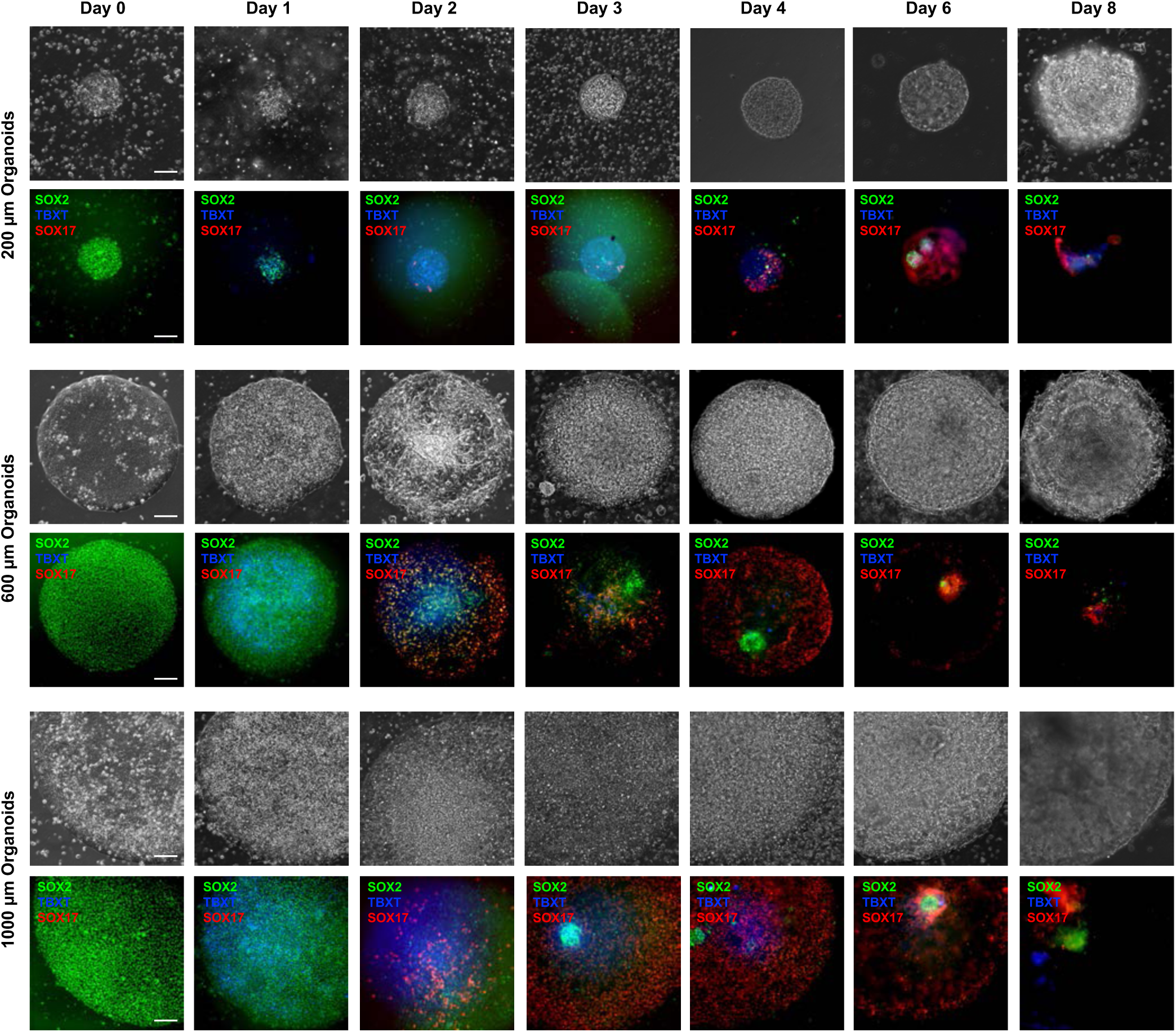
Cell fate transition during mesoderm-endoderm divergence (related to Figure 4). Representative images of brightfield morphology and fluorescence signals for SOX2–mCitrine, BRA–mCerulean, and SOX17– tdTomato in micropatterned RUES2-GLR hESCs with different sizes across different time points (Day 0–Day 8). The images illustrate the decline of SOX2 expression, with TBXT (BRA) and SOX17 emerging at Day 1 and peaking at Day 2.

**Figure S6.**
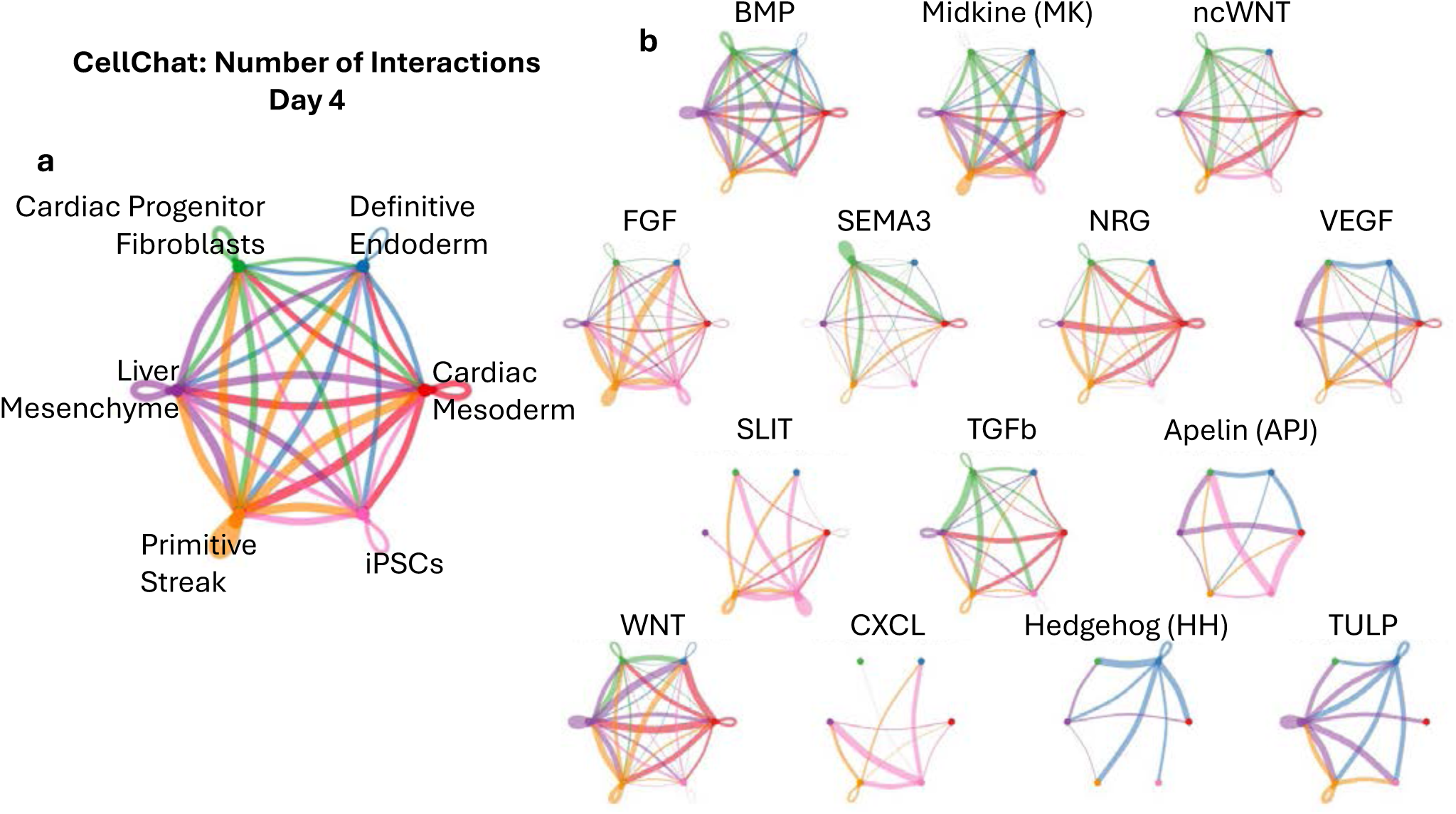
Ligand-receptor analysis on Day 4 during mesoderm-endoderm divergence (related to Figure 4). (a) Cell-cell interaction networks for all detected signaling pathways. (b) Cell-cell interactions highlighting selected embryogenesis-related signaling pathways.

**Figure S7.**
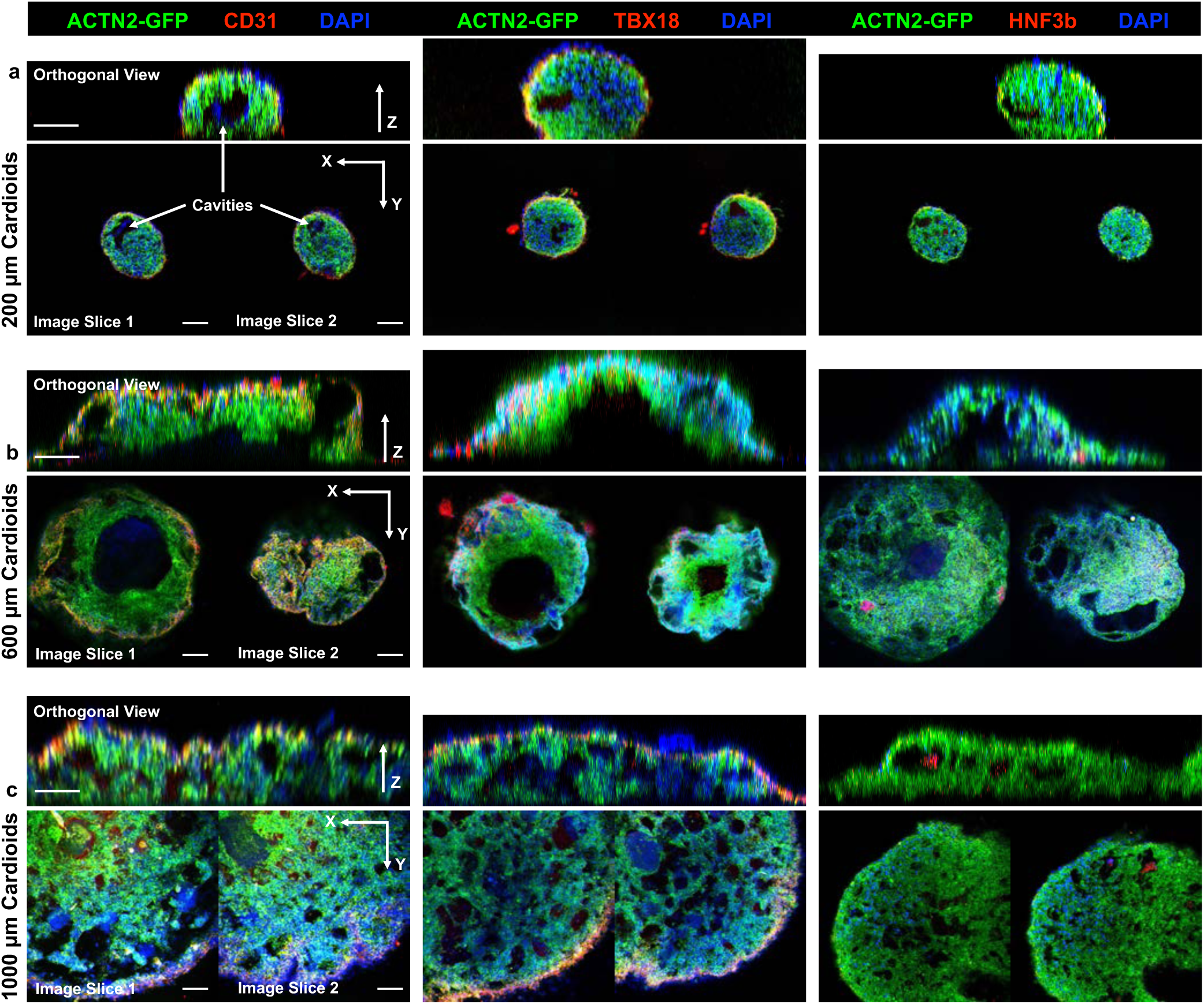
Cavitation inside Day 21 cardioids generated from different pattern sizes (related to Figure 5). Representative two-photon microscopy images of micropatterned cardioids generated from ACTN2-GFP reporter hiPSC line and stained with endothelial marker (CD31), epicardial marker (TBX18), and foregut marker (HNF3b). 3D reconstructed orthogonal sideview of entire whole-mounted cardioids (upper Z-direction) and two single-plane image slices in the middle of each cardioids (bottom two). (a) 200 µm cardioids showed limited cavitation with isolated small cavities. (b) 600 µm cardioids showed large cavity formation resembling early heart chamber. (c) 1000 µm cardioids showed interconnected mesh-like cavitation. Scale bar: 100 µm.

**Figure S8.**
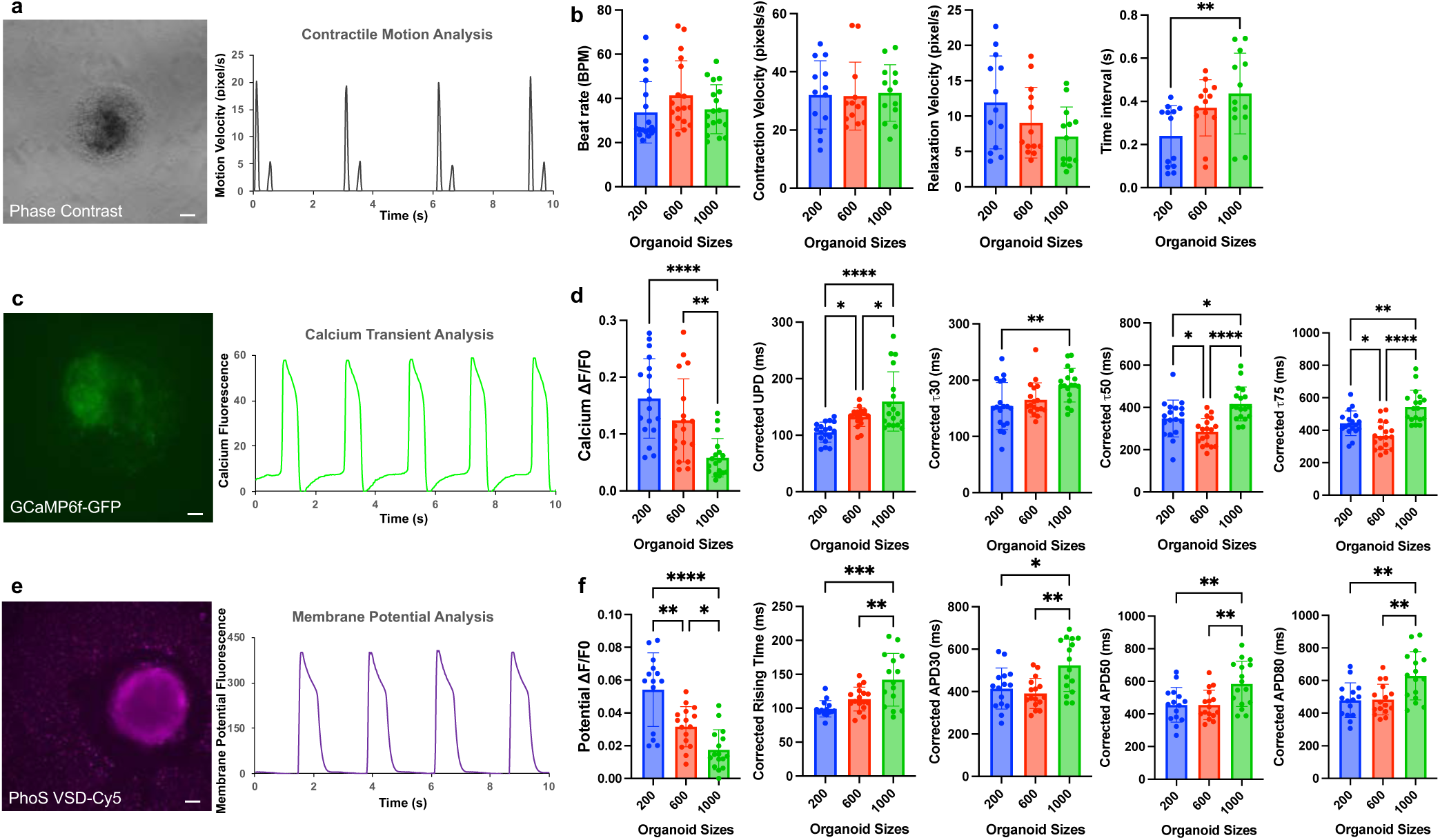
Contractile function analysis of Day 21 micropatterned cardioids with different sizes (related to Figure 5). (a) Representative brightfield image (600 µm cardioids) used for contractile motion analysis, revealing (b) an elongated contraction-relaxation time interval in larger 1000 µm cardioids. (c) Representative GCaMP6f-GFP fluorescence image (600 µm cardioids) for calcium transient analysis. (d) 600 µm cardioids exhibited the shortest calcium decay time, whereas 1000 µm cardioids displayed prolonged calcium transients. (e) Representative PhoS VSD-Cy5 fluorescence image (600 µm cardioids) for membrane potential analysis, indicating that 1000 µm cardioids had an extended action potential duration. Scale bar: 50 µm. Sample size > 10 cardioids from three independent differentiation. Statistical analysis: one-way ANOVA with post-hoc Tukey analysis (*p < 0.05, **p<0.01, ***p<0.001, ****p<0.001).

**Figure S9.**
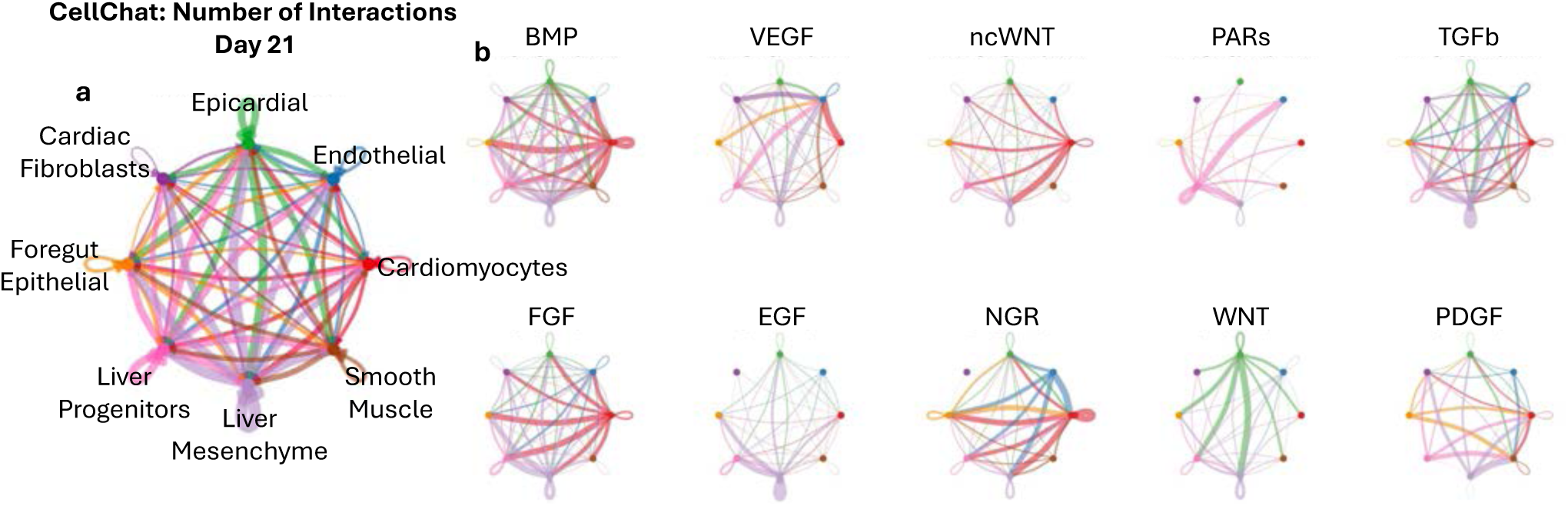
Ligand-receptor analysis on Day 21 for heart-foregut crosstalk (related to Figure 5). (a) Cell-cell interaction networks for all detected signaling pathways. (b) Cell-cell interactions highlighting selected embryogenesis-related signaling pathways.

**Table S1:** Antibodies used in the study.

| **Table S1: Antibodies used in the study** |  |  |
| --- | --- | --- |
| ***Primary antibody*** |  |  |
| CDH5 | V1514-200ul | Sigma |
| PECAM1 | PA5-32321 | Invitrogen |
| HNF3B | sc-374376 | Santa Cruz |
| HNF3A | ab170933 | Abcam |
| TBX18 | sc-130428 | Santa Cruz |
| WT1 | PA5-16879 | Invitrogen |
| TNNI | ab47003 | Abcam |
| TNNT2 | MA5-12960 | Invitrogen |
| BeRST 1 | PhoS1-10 | PhotoSwitch Bioscience |
| ***Secondary Antibody*** |  |  |
| Goat anti-Mouse IgG, Alexa Fluor 488 | A-11029 | Invitrogen |
| Goat anti-Rabbit IgG, Alexa Fluor 488 | A11008 | Invitrogen |
| Donkey anti-Goat IgG, Alexa Fluor 546 | A11056 | Invitrogen |
| Goat anti-Mouse IgG, Alexa Fluor 546 | A11003 | Invitrogen |
| Goat anti-Rabbit IgG, Alexa Fluor 546 | A11010 | Invitrogen |

## Supplemental Movies

**Movie S1.** 3D reconstructed confocal microscopy image of a micropatterned cardioid stained with cardiac troponin (TNNI2) and vimentin (VIM).

**Movie S2.** Cross-sectioning of 3D reconstructed two-photon microscopy image of a micropatterned cardioid generated from TNNI2-GFP hiPSC reporter line and stained with endodermal marker HNF3A.

**Movie S3.** Brightfield video of a beating 600 µm cardioid.

**Movie S4.** Fluorescent video of a beating 600 µm cardioid generated from GCaMP6f-GFP reporter hiPSC line for calcium transient analysis.

**Movie S5.** Fluorescent video of a beating 600 µm cardioid stained by membrane potential dye (PhoS VSD-Cy5) for action potential analysis.

